# Impact of Microbial Iron Oxide Reduction on the Transport of Diffusible Tracers and Non-diffusible Nanoparticles in Soils

**DOI:** 10.1101/386128

**Authors:** Xiaolong Liang, Mark Radosevich, Frank Löffler, Sean M. Schaeffer, Jie Zhuang

## Abstract

**Abstract:** *In situ* bioremediation to achieve immobilization of toxic metals and radionuclides or detoxification of chlorinated solvents relies on electron donor additions. This practice promotes microbial Fe(III)-oxide mineral reduction that could change soil pore structure, release soil colloids, alter matrix surface properties, and cause the formation of secondary (i.e., reduced) Fe-mineral phases. These processes in turn may impact rates of bioremediation, groundwater quality, and ultimately contaminant fate. Continuous flow columns packed with water-stable soil aggregates high in Fe-oxides were infused with artificial groundwater containing acetate as electron donor and operated for 20 or 60 days inside an anoxic chamber. Soluble Fe(II) and soil colloids were detected in the effluent within one week after initiation of the acetate addition, demonstrating Fe(III)-bioreduction and colloid formation. Br^-^, 2,6-difluorobenzoate (DFBA), and silica-shelled silver nanoparticles (SSSNP) were selected as diffusible tracer, low-diffusible tracer, and non-diffusible nanoparticles, respectively, to perform transport experiments before and after the active 20-day bioreduction phase, with an aim of assessing the changes in soil structure and surface chemical properties resulting from Fe(III)-bioreduction. The transport of diffusible Br^-^ was not influenced by the Fe(III)-bioreduction as evidenced by identical breakthrough curves before and after the introduction of acetate. Low-diffusible DFBA showed earlier breakthrough and less tailing after the bioreduction, suggesting alterations in flow paths and surface chemical properties of the soils. Similarly, non-diffusible SSSNP exhibited early breakthrough and enhanced transport after the bioreduction phase. Unexpectedly, the bioreduction caused complete retention of SSSNP in the soil columns when the acetate injection was extended from 20 days to 60 days, though no changes were observed for Br^-^ and DFBA during the extended bioreduction period. The large change in the transport of SSSNP was attributed to the enhancement of soil aggregate breakdown and soil colloid release causing mechanical straining of SSSNP and the exposure of iron oxide surfaces previously unavailable within aggregate interiors favorable to the attachment of SSSNP. These results demonstrate that microbial activity can affect soil properties and transport behaviors of diffusivity-varying solutes and colloids in a time dependent fashion, a finding with implication for interpreting the data generated from soil column experiments under continuous flow.

**Highlights:**

- Fe(III)-bioreduction causes time-dependent aggregate breakdown and colloid release.
- Short-term bioreduction alters soil aggregate surface chemistry and tracer transport.
- Electron donor amendment enhances transport of nanoparticle tracer.

## 1. Introduction

Subsurface bioremediation brought about by electron donor addition creates anoxic conditions that stimulate the growth of iron reducing and/or sulfate reducing bacteria (Chapelle and Lovley, 1992; Si et al., 2015). Subsequently, oxidized forms of iron are reduced to Fe(II), solubilizing iron oxide minerals. The soluble Fe(II) and secondary precipitation of iron (e.g., ferrous hydroxide) can result in abiotic transformation of contaminants and/or the release of colloidal clay and iron mineral colloids (Pedersen et al., 2006; Thompson et al., 2006). Adsorption of contaminants to these colloids may enhance contaminant transport via colloid-facilitated transport (Bose and Sharma, 2002; Zhuang et al., 2003). Additionally, biomass increase and colloid production can cause pore clogging, potentially reducing hydraulic conductivity of the porous medium down gradient from the treatment zone. Alternatively, advective flow paths and increased hydraulic conductivity may trigger substantial soil aggregate breakdown. Thus, altering the indigenous properties of subsurface media may impact coupled processes controlling the fate and transport of contaminants, and cause unintended secondary impacts on the properties of the porous media and groundwater quality (e.g. secondary mineral precipitates, permeability, and microbial activity).

Anaerobic bioremediation is an attractive technology for subsurface soil and water remediation based on cost and effectiveness (Ellis et al., 2000; Coates and Anderson, 2000; Liang et al., 2017). The technology generally aims to create anoxic conditions via addition of soluble electron donors, such as acetate and lactate or higher molecular weight substrates, for stimulating microorganisms that degrade organic contaminants (e.g. chlorinated solvents) and reduce heavy metals and radionuclides to insoluble forms thereby immobilizing them *in situ* (Aulenta et al., 2006). Once anoxic conditions are achieved, anaerobic respiration with available electron acceptors is stimulated leading to biologically-mediated reduction of Fe(III)-oxide minerals (the most common mineral oxide of soils and subsurface environments) and the formation of soluble Fe(II) (Caldwell et al., 1999; Weber et al., 2006; Mejia et al., 2016). Iron oxides, which have diverse crystallinities and reactivity (e.g. ferrihydrite, goethite, lepidocrocite, and hematite), are extensively present in soils (Pedersen et al., 2006; Vink et al., 2017). The indigenous iron oxides serve a very important role as aggregating agents that “cement” clay particles together into aggregates (Goldberg et al., 1990; Braunschweig et al., 2013). Reduction of Fe(III)-oxides under anoxic conditions may cause disintegration of soil aggregates and generate mobile colloids (Hansel et al., 2005; De-Campos et al., 2009). These processes may disrupt pore structure and alter pore connectivity, flow paths, and permeability (Guan et al., 2017). These alterations could either promote or inhibit the transport of solutes and colloids. Schaider et al. (2014) found that iron oxide aggregates can alter the transport of particulate particles and sequester metals. The formation and mobilization of colloids can act as vectors to facilitate co-transport of solutes and toxic metals in soils and groundwater (McCarthy and McKay, 2004; Maurice and Hocella, 2008; Guan et al., 2017). The reduction of Fe(III)-oxides may also change soil surface properties to influence the reactive transport processes (Hansel et al., 2003; Jardine, 2008).

Transport of solutes and colloids in soil is influenced by the pore structure and surface chemistry of soils, solution chemistry, and hydrological conditions (Zhuang et al., 2005; 2007; 2010; Bradford and Torkzaban, 2008; Mohanty et al., 2016; Pachapur et al., 2016). The influencing mechanisms have been well examined at varying scales from laboratory columns (repacked or undisturbed) to field scale (McKay et al., 2000; Arora et al., 2015; Karadimitriou et al., 2017). Bioremediation treatment may alter aquifer porosity, flow paths, and mineral interfacial properties and in turn change the attenuation and migration of solutes, colloids, and microbial cells; all these may exert feedback effects on microbial bioremediation. Microorganisms, nutrients, or electron donors are generally applied to accelerate in situ bioremediation (Ellis et al., 2000; Lovley 2003; Moon et al., 2017), yet the remediation efficiency is subject to their mobility in porous media (Song et al., 2017). Thus far, few studies have addressed the impacts of biostimulation on the transport of solutes and colloids, making difficult to resolve the low-efficiency problem of bioremediation under field conditions. Therefore, in-depth investigations are needed to understand the potential that biostimulation influences the mobility of solutes and colloids including microorganisms.

The objective of this research was to assess the impact of biologically-mediated Fe(III)-oxide reduction on the transport of solutes and colloids with respect to soil structure breakdown under saturated flow conditions, shedding light on the interplays of microbial activities with the solutes and colloids migration during bioremediation processes. Breakthrough tests with diffusivity-varying tracers and non-diffusing nanoparticles both before and after acetate-stimulated Fe(III)-bioreduction were conducted to evaluate the alteration of soil surface chemistry and flow pathways. The research provides significant insights into the feedback effects of anoxic bioremediation on the transport of solutes and colloids, microbial distribution, and soil aggregate structure.

## 2. Materials and methods

### 2.1 Porous media

The columns were packed with different porous media, including uncoated and goethite-coated silica sand and water-stable soil aggregates extracted from an iron oxide-rich natural soil. The sand grains had a median diameter (*d*_50_) of 0.25 ± 0.01 mm with a trade name Accusand (Grade 50/70, Unimin Corporation, New Canaan, CT, USA). Prior to coating with goethite, the sand was chemically treated to remove natural metal oxides from the grains following the established procedure (Zhuang and Jin, 2003). Goethite synthesis and coating on the cleaned sand were performed as described by Zhuang and Jin (2008). The natural soil was collected from an eroded agricultural site mapped as the Decatur silty clay loam. The Decatur series is a fine, kaolinitic, thermic rhodic Paleudults. Soil aggregates were extracted by wet sieving of the bulk soils through 2,000, 250, and 53 μm sieves using the modified method as described in Zhuang et al. (2008). Two fractions of water-stable soil aggregates, microaggregates (53-250 μm) and macroaggregates (250-2000 μm), were obtained and then air-dried for experimental use. The citrate-bicarbonate-dithionite extractable iron (Mehra and Jackson, 1960) of the bulk soil, microaggregates, and macroaggregates were 5.5%, 4.7% and 5.2% (w/w), respectively, as measured using the ferrozine method (Viollier et al., 2000).

### 2.2 Tracers and nanoparticles

Two diffusible tracers and one non-diffusing nanoparticle were used for transport experiments, including bromide (Br^-^ in KBr) (ionic diffusible tracer), 2,6-difluorobenzoate (DFBA) (molecular diffusible tracer with lower diffusivity than Br^-^) (Mayes et al., 2003), and silica-shelled silver nanoparticles (SSSNP) (non-diffusible particle). The SSSNP was purchased from nanoComposix (http://nanocomposix.com/) and has a core of silver nanoparticles with average diameter of 106 nm. The silver cores are encased in a shell of silica with average thickness of 22 nm, resulting in a total particle diameter of 150 nm. The SSSNP were negatively charged, with a measured zeta potential of -5.7 mV at pH 8. SSSNP were specifically selected as a “non-diffusing” particle and potentially as a non-reactive particle given the silica shell surrounding the silver-core. Advantages to using these shelled particles instead of other previously used particle tracers, such as viruses (e.g., MS-2), are their resistance to biotic and abiotic breakdown, the ease and accuracy of quantification in the effluent samples using graphite furnace atomic absorption spectroscopy, and the convenience to distinguish introduced SSSNP from native soil colloids for mechanistic understanding of transport processes.

### 2.3 Bacterial strain, growth media, and inoculation for stimulated bioreduction

*Geobacter* species, with capacity to oxidize organic compounds coupled with reduction of iron oxides or other metal minerals, are ubiquitous in subsurface environments (Caccavo et al., 1994). As such, *Geobacter*, and other microorganisms with similar metabolism have been extensively studied and used for anaerobic bioremediation (Lovley 2003; Moon et al., 2017). To ensure active Fe(III) bioreduction, the soil macroaggregates (250-2,000 μm) were inoculated with laboratory-grown culture of *Geobacter sulfurreducens* strain PCA (ATCC 51573; Caccavo et al., 1994) prior to packing the columns.

Specifically, *G. sulfurreducens* was grown in mineral salts medium containing 1.0 g of NaCl, 0. 5 g of MgCl_2_, 0.2 g of KH_2_PO_4_, 0.3 g of NH_4_Cl, 0.3 g of KCl, 0.015 g of CaCl_2_, 1 mg of resazurin, and 2 ml of trace element solution, amended with 5 mM acetate and 10 mM ferric citrate per liter was prepared as described by Löffler et al. (1996). The prepared medium was boiled and transferred to serum bottles while flushing with oxygen-free 80/20 (v/v) N_2_-CO_2_, and the pH was adjusted to 7.2 with flow of CO2. The serum bottles were autoclaved, and filter-sterilized (with a 0.22 μm Millex filter syringe) acetate and ferric citrate were added to the medium to a final concentration of with 5 mM and 10 mM, separately. The serum bottles inoculated with *G. sulfurreducens* were cultured at 30 °C (Löffler et al., 1996). Once the cultures reached stationary phase (3-5 d; optical density at 600 nm = 0.2 to 0.35; cell concentration of approximately 1 × 10^8^ cells per milliliter), 200 ml of culture suspension was centrifuged at 4,248 g, and the bacterial pellet was resuspended in 15 ml of the growth medium in the anaerobic chamber with an 80/20 (v/v) N_2_/CO_2_ atmosphere. Then, all the resuspended *G. sulfureducens* were uniformly sprayed onto 600 g of air-dried but non-sterile, water-stable soil macroaggregates that were thinly spread on a tray inside the anaerobic chamber. During the application of cell suspension, the aggregates were continuously mixed using a glass rod to achieve uniform inoculation of the bacteria.

### 2.4 Transport experiment

All transport experiments were conducted in plexiglass (acrylic) columns (25 cm in length with an inside diameter of 3.8 cm) with input solution introduced from the bottom of the column in pulse input mode through a peristaltic pump at pore velocity of 24.4 cm/h. Teflon tubing was used throughout the system except for a portion of tygon tubing needed in the pump. The columns were fitted with five ports connected to pressure sensors (Honeywell Sensing and Control, Inc., USA), which were separated by 5-cm intervals along the column length. Real-time data of hydrostatic pressure were collected with data loggers of CR-1000 Measurement and Control Systems (Campbell Scientific, Inc., Logan Utah) to calculate hydraulic conductivity between different sections of the columns according to the difference in pressure. During the transport experiment, liquid effluent was collected from the top of the column into 20-mL glass tubes using Retriever II fraction collectors for determining the concentrations of tracers, iron, and or colloids as described in section 2.5. The protocols for column experiments are shown in supplementary materials (Table S1 and Fig. S1).

Three sets of separate transport experiments were conducted using vertical columns under saturated steady-state flow conditions. The first set aimed to evaluate the appropriateness of use of SSSNP as non-diffusible particle tracer and the effect of iron oxide on the transport of tracers and nanoparticles. The experiments included two columns that were wet-packed with uncoated and goethite-coated sands, respectively. The sand columns were flushed with KCl solution (0.67 mM, pH 6.5) prior to the tracer experiments. The input solution for the sand columns contained Br^-^ (50 mg L^-1^ KBr), DFBA (40 mg/L), and SSSNP (40 ug L^-1^) in the KCl solution.

The second set of experiments aimed to evaluate the effect of Fe(III)-bioreduction on the transport of tracers using five columns dry-packed with *Geobactor*-inoculated soil macroaggregates under anoxic conditions. The experiments included three acetate-stimulated Fe(III)-bioreduction columns (one with 20 days of continuous injection of acetate and two replicates with 60 days of acetate injection) and two control columns (no acetate addition). The 60-day experiments aimed to corroborate the results of bioreduction effects observed from the 20-day experiments. After dry packing, the soil aggregate columns were flushed with carbon dioxide to replace the air in soil pores, followed by flushing with KCl solution (0.67 mM, pH 6.5) to achieve fully saturated conditions without remaining gas pockets. Each column experiment consisted of three phases with constant level of total ionic strength of solutions (2 mM). Before bioreduction, transport experiments with the KCl input solution containing Br^-^ (85 mg/L KBr), DFBA (50 mg/L), and SSSNP (40 μg/L) were performed in all columns (phase 1). In bioreduction process (phase 2), the columns were flushed with artificial groundwater solution (AGW), which had a total ionic strength of 2 mM and a pH value of 7.5, consisting of CaCl_2_ (0.075 mM), MgCl_2_ (0.082 mM), KCl (0.051 mM), and NaHCO_3_ (1.5 mM), modified from Ferris et al. (2004). The AGW contained trace elements, vitamins, and acetate (bioreduction-stimulated column) or without acetate (control column) (Wolin et al., 1963). Acetate added to the columns served as the electron donor for *Geobacter* Effluent samples from columns during bioreduction were analyzed for the concentrations of Fe(II), Fe(III) and colloids as described in Section 2.5. After 20 or 60 days of bioreduction, the same transport experiment as that prior to bioreduction was performed (phase 3). Breakthrough and elution data for Br^-^, DFBA, and SSSNP were collected during phases 1 and 3. At the conclusion of the above procedures, the columns were sectioned in 5-cm intervals along the longitudinal flow path of the columns. The distribution of *Geobacter* and readily deducible iron content in soil aggregates from each section were investigated as described in Section 2.7 and 2.8.

The third set of experiments was conducted to examine the effects of aggregate size fractions on the transport of tracers and nanoparticles outside the anoxic chamber without bioreduction treatment (exposed to oxygen), since exposure to aerobic conditions can suppress reduction of iron oxides. The experiments included two columns, which were dry packed with microaggregates and macroaggregates, respectively. The experimental procedures were the same as those used in the second set of column experiments.

### 2.5 Chemical analysis

Bromide concentrations in the effluent fractions were determined using ion chromatography as described elsewhere (Qin et al., 2017). The concentration of DFBA was measured with a modified HPLC method (Galdiga and Greibrokk, 1998). Briefly, DFBA was resolved from other effluent constituents using an Econosphere C-18 RP column (5 μm, 150 mm x 4.6 mm) with isocratic elution using a mobile phase consisting of 95% K-phosphate buffer (5 mM, pH 3) and 5% acetonitrile (v/v) at a flow rate of 1.0 mL min^-1^. DFBA was quantified using UV absorption at 200 nm and the concentration was calculated via linear regression of peak area of external DFBA standard solutions over a concentration range from 1 to 50 mg L^-1^. All samples were diluted 1:10 (v/v) in mobile phase to minimize sample matrix effects and filtered through 0.1 μm membrane filters (Merck Millipore Ltd., Cork, Ireland) prior to analysis. SSSNP (with hydrodynamic diameter of 129.8 nm) in effluent fractions were quantified by measuring the concentration of silver by graphite furnace atomic absorption spectrometry using a Perkin-Elmer Graphite Furnance AA equipped with a transversely heated graphite atomizer as described by Fernández et al. (2010). Effluent samples were diluted 10^4^ times with deionized water before analysis, and 20 μL of diluted sample was injected with 10 μL of matrix modifier (prepared by dissolving 0.05 mg de Pd and 0.003 mg Mg(NO_3_)_2_ in 10 μL 1% HNO_3_). The silver detection program in furnance AA was described in Fernández et al. (2010).

### 2.6 Numerical modeling

The HYDRUS code (Šimůnek et al., 2008) simulating saturated water flow based on the Richards equation was used to simulate the transport of bromide, DFBA, and SSSNP. Transport behaviors of Br^-^ and 2,6-DFBA were simulated using the classical advection-dispersion equation (ADE). The transport and retention of SSSNP were simulated using the ADE with first-order terms for kinetic retention and release as described in the HYDRUS code. The equation is given in modeling the transport behavior as:

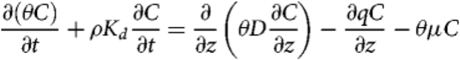

where θ[L^3^L^−3^] is the pore volume in the column, C [M L^−3^; M represents the units of mass] is the concentration of bromide, DFBA, and SSSNP in the aqueous phase, ρ [M L^−3^] is the bulk density of soil aggregates, K_d_ [L^3^ M^−1^] is the adsorption coefficient to soil aggregates, t is time [T, T represents time units], D [L^2^ T^−1^] is the hydrodynamic dispersion coefficient, z [L] is the distance from the inlet of a column, q [L T^−1^] is the Darcy velocity of input solutions, and μ [T^−1^] is the first-order retention coefficient for tracer transformation processes.

### 2.7 Assessment of soil aggregate properties

Component analysis of soil aggregates were performed by commercial service of Midwest Laboratories Inc. (Omaha, NE, USA). Particle size distribution of soil aggregates was analyzed to indicate the reactivity behaviors of soil aggregates. The procedures and principles were described in Kemper and Rosenau (1986). Fifty grams of air-dried soil aggregates were presoaked in distilled water for 30 min and sieved with a top-down sequence of seven sieves of 2,000, 840, 300, 250, 150, 90, and 53 mm mesh size. The sieves with the contents were oscillated vertically in water with an amplitude of 4 cm at a rate of one oscillation per second for twenty times. The retained aggregates on each sieve after wet-sieving were recovered and dried at 50 °C in a drying oven. The particle size distribution was calculated with dry weight fractions. After bioreduction, dispersion of the aggregates was assessed based on readily reducible iron content. Soil aggregates from each section were homogenized, and 5-g sub-samples were placed in 50-mL centrifuge tubes, in which soil aggregates were shaken vigorously with 40-mL distilled water in an ice-water bath for 30 min (Kemper and Rosenau, 1986; Viollier et al., 2000). The mixtures were centrifuged at 4,248 g for 20 min, and the readily reducible iron in supernatant were determined using a modified Ferrozine method (Stookey, 1970). During column bioreduction experiments, the concentrations of Fe(II) and Fe(III) in the effluent were measured using a modified Ferrozine method (Stookey, 1970). Colloids were determined by centrifuging the effluent sample at 4,248 g for 20 min and measuring the mass of pellets after oven-drying, assuming the mass of dissolved salts was negligible.

### 2.8 DNA extraction and sequencing

To analyze the distribution of *Geobacter* in soil aggregates, DNA was extracted from fractioned soil aggregates using PowerLyser PowerSoil DNA isolation kit (MoBio Laboratories Inv. Carlsbad, California, USA). The DNA samples were quantified using PicoGreen Assay Kit (Carlsbad, CA, USA) and sent out for sequencing at HudsonAlpha Genomics Services Lab (Huntsville, AL, USA). The V3-V4 region of 16S rRNA gene of bacteria were amplified with primers of 341F_CCTACG GGNGGCWGCAG and 785R_GACTACHVGGGTATCTAATCC) in PCR. Finally, sequencing was performed using 300PE (paired-end) on the Illumina MiSeq platform (Illumina, USA). All methods were performed according to the manufacturers’ protocol. Sequences analyses were performed using MOTHUR per standard operating procedure (Kozich et al., 2013). The results of these sequence analyses were used to calculate the relative abundance of *Geobacter* along the flow path in the columns.

## 3 Results and discussion

### 3.1 Effect of iron oxide on transport

The uncoated and goethite-coated quartz sands were used as simple and stable porous systems to evaluate the effect of iron oxide on the transport of tracers and nanoparticles. Br^-^ was included in the input solution to quantify transport behaviors of a diffusible tracer and to evaluate uniformity and integrity of the column packing in terms of hydrodynamic dispersion. As a conservative tracer, Br^-^ showed ideal and complete transport behavior in the sand with and without goethite coating (Fig. 1). In comparison, the breakthrough of SSSNP from the uncoated sand was slightly retarded relative to that of Br^-^ and eventually reached a stable maximum relative concentration (max C/C0) of 0.85. However, almost no SSSNP broke through the goethite-coated sand column. The fitted parameters of the breakthrough curves of SSSNP showed a 14-fold increase in attachment and a 3-fold decrease in detachment of SSSNP in goethite-coated sand compared with the uncoated sand columns. The maximum solid phase concentration of SSSNP was ~20 times higher in the goethite-coated sand than in the uncoated sand, suggesting a strong affinity of the SSSNP to the iron-oxide surface (Table 1). The strong attachment of SSSNP to goethite-coated sand in this study was consistent with previous results showing sequestration of silver nanoparticle (no silica-shell) by iron oxides (Sagee et al., 2012; Liang et al., 2013).

**Fig. 1.**
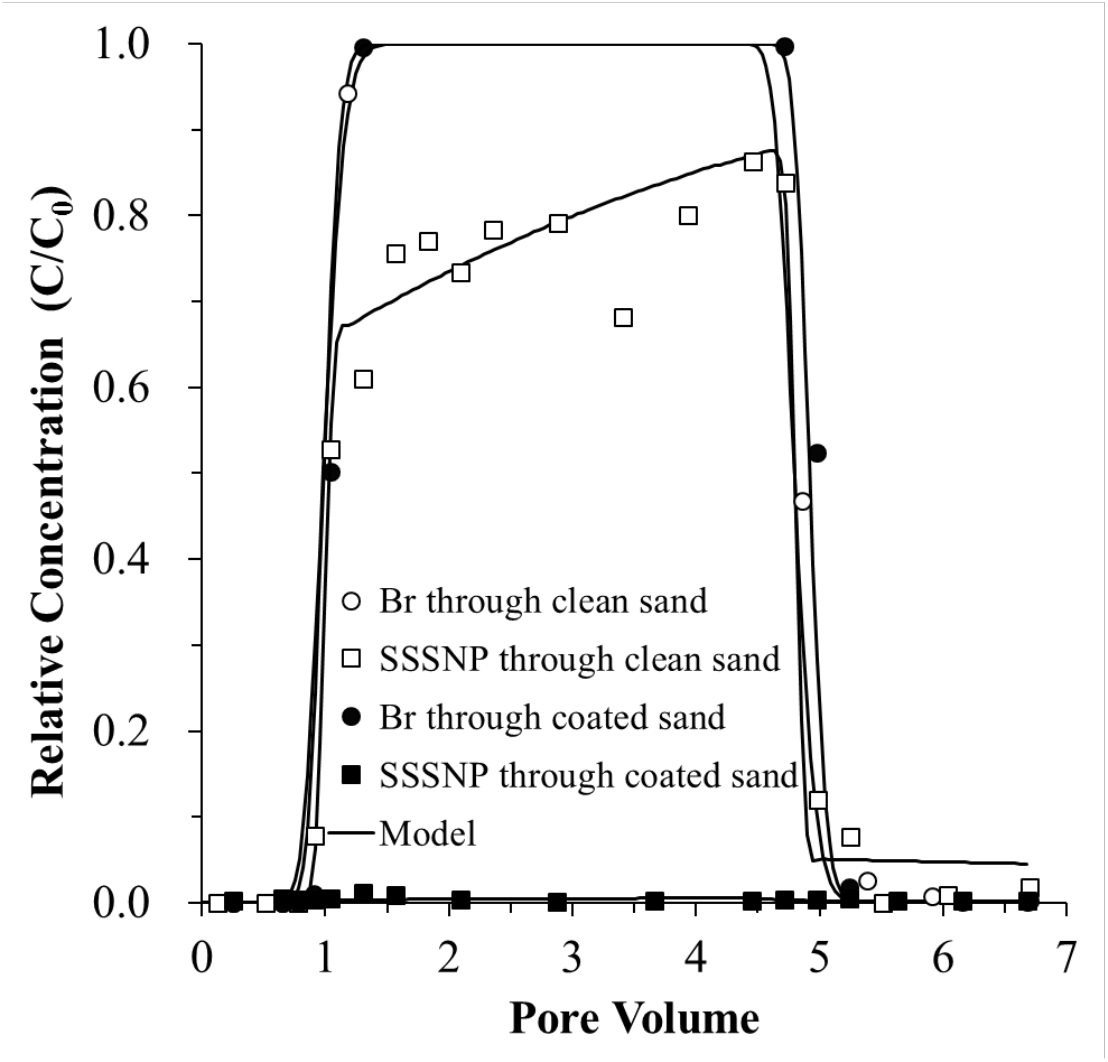
Transport of bromide and silica-shelled silver nanoparticles (SSSNP) through goethite-coated and uncoated sands. The concentrations of bromide and SSSNP are shown over pore volume with columns flushed using KCl solution (0.67 mM, pH 6.5).

**Table 1.**
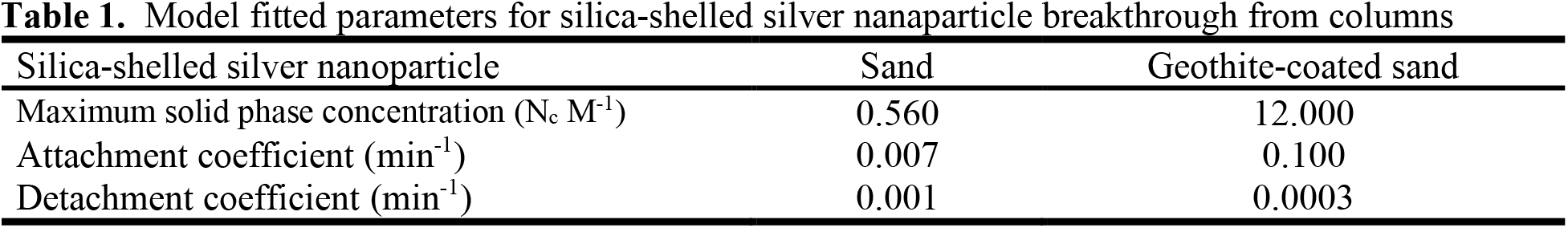
Model fitted parameters for silica-shelled silver nanaparticle breakthrough from columns

| Silica-shelled silver nanoparticle | Sand | Geothite-coated sand |
| --- | --- | --- |
| Maximum solid phase concentration ($N_c \text{ M}^{-1}$) | 0.560 | 12.000 |
| Attachment coefficient ($\text{min}^{-1}$) | 0.007 | 0.100 |
| Detachment coefficient ($\text{min}^{-1}$) | 0.001 | 0.0003 |

### 3.2 Effect of aggregates on transport

To further determine if the change in soil aggregate sizes primarily influences the transport of DFBA and SSSNP in column experiments, transport experiments were conducted in columns packed with water-stable macroaggregates (250-2,000 μm) and microaggregates (53-250 μm) under oxic conditions and without any reduction of Fe(III) oxides. The transport of bromide through both aggregate fractions showed no obvious differences (Fig. 2); however, DFBA was retarded in both columns, with larger retardation in the microaggregates (K_d_ of 0.12) than in the macroaggregates (K_d_ of 0.06). Since the surface properties of macroaggregates and microaggregates should have been very similar, the larger retardation is ascribed to the greater total surface area and smaller pores of the microaggregates compared to the macroaggregates. The breakthrough of SSSNP was negligible with both columns likely due to the strong interactions of SSSNP with Fe(III) oxides (Fig. 2). Similar results of decreased silver nanoparticle mobility in smaller soil aggregates were reported by Sagee et al. (2012), in which the explanation was proposed that the reaction of silver nanoparticles with soil occurs at the aggregate surface, and the smaller aggregates have increased surface area. Aggregate structure has been shown to play an important role in retention of large molecules and nanoparticles with complicated interacting environmental factors (Sagee et al., 2012; Liang et al., 2013). For example, transport of silver nanoparticles in natural soil showed that aggregate size exerted major influence on silver nanoparticle transport, with silver nanoparticle mobility increased in the column of larger soil aggregates (Sagee et al., 2012). The transport and retention of silver nanoparticles in sand columns by Liang et al. (2013) also found that the mobility of silver nanoparticles was enhanced by increase in sand grain size. The soil aggregates used in this study were highly rich in iron oxides and had strong binding capacity for SSSNP, which caused complete retention of SSSNP in both microaggregates and macroaggregates columns. In contrast, the retardation of DFBA increased in the microaggregates.

**Fig. 2.**
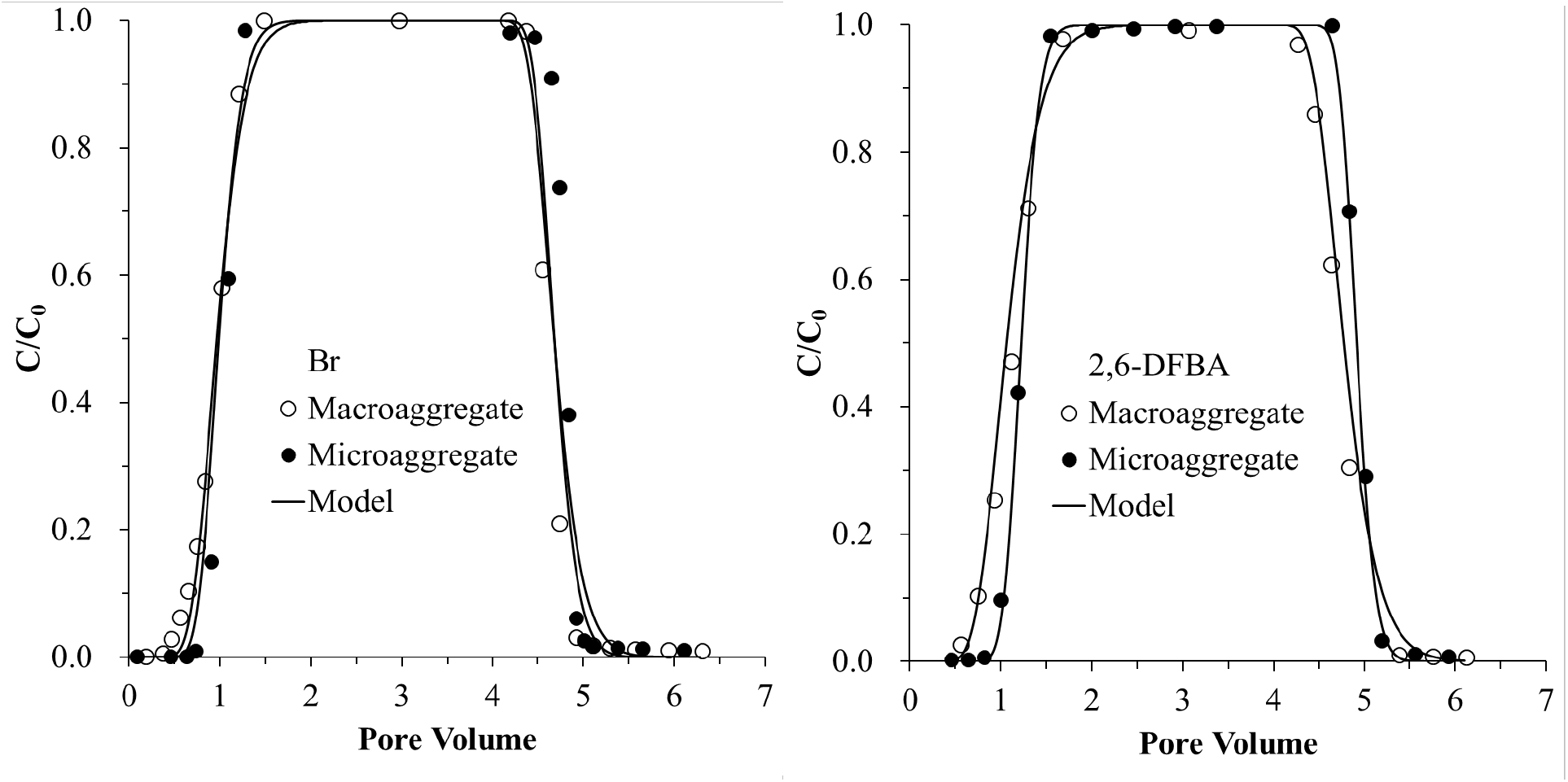
Transport of bromide and DFBA through columns packed with water-stable macroaggregaes (250-2,000 μm) or water-stable microaggregates (53-250 μm) under oxic conditions without Fe(III)-bioreduction.

### 3.3 Effect of short time bioreduction on transport

Soil aggregates rich in iron oxides were used to examine the effect of Fe(III)-bioreduction on transport behaviors of tracers and nanoparticles. The breakthrough of Br^-^ occurred at approximately one pore volume before and after the Fe(III)-bioreduction phase (Fig. 3), indicating that Fe(III)-bioreduction did not influence the transport of the conservative tracer. Transport of DFBA exhibited some retardation and tailing in the breakthrough experiment before the bioreduction, but the retardation was eliminated after the bioreduction phase (Figs. 3 and 4). This result suggests that Fe(III)-bioreduction induced either physical and/or chemical changes of the aggregates. Modeling results showed that the estimated dispersivity (*D*) of DFBA during transport remained similar before and after the Fe(III)-bioreduction phase while the estimated DFBA sorption coefficient (*K_d_*) was about one order of magnitude lower after Fe(III)-bioreduction (Table 2). SSSNP exhibited a very pronounced response to the Fe(III)-bioreduction treatment. Almost no breakthrough of SSSNP was observed before Fe(III)-bioreduction (i.e., in phase 1), whereas the relative concentrations (C/C0) of SSSNP in the effluent reached only 0.3 after the Fe(III)-bioreduction (i.e., in phase 3) (Fig. 3). The estimated maximum solid phase concentration of SSSNP in phase 3 was one fourth that in phase 1 with a lower attachment coefficient. The estimated detachment coefficient of SSSNP was 100-fold greater in phase 3 than in phase 1 (Table 2). These results indicate that the presence of Fe(III) oxides greatly reduced the mobility of the nanoparticle tracer. Ryan et al. (1999) found that the bacteriophage particles and silica colloids attached to iron oxide-coated sand could be mobilized by anionic surfactant, elevated pH, and reductant, suggesting that iron oxide removal could promote the detachment of colloids and bacteriophage. Vink et al. (2017) showed that arsenic release corresponded to the fractions of readily reducible iron in sediments. At the initial phase of incubation with acetate-supplemented AGW solution, microbially mediated bioreduction released relatively small amount of Fe(II) from soil aggregates, causing certain alteration of the aggregate surface properties. Since the binding capacity for metals and colloids are highly dependent on ferric phases in soils (Pedersen et al., 2006), the reduction of iron oxides can lead to release of certain amount of retained molecules and colloids, such as DFBA and SSSNP.

**Fig. 3.**
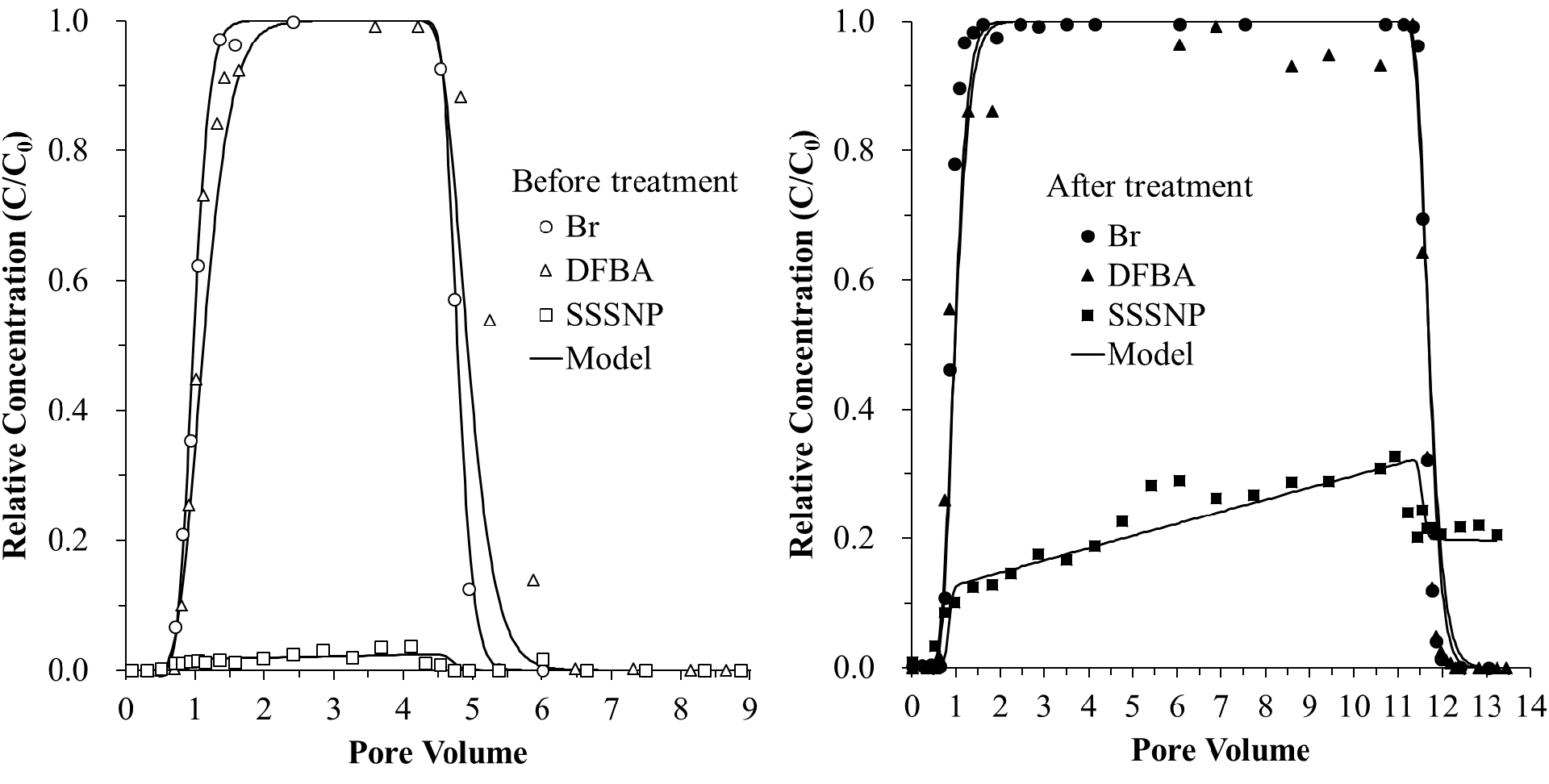
Breakthrough curves of tracers (Br^-^ and DFBA) and silica-shelled silver nanoparticles (SSSNP) from columns packed with water-stable soil aggregates before and after the 20-day Fe(III)-bioreduction.

**Fig. 4.**
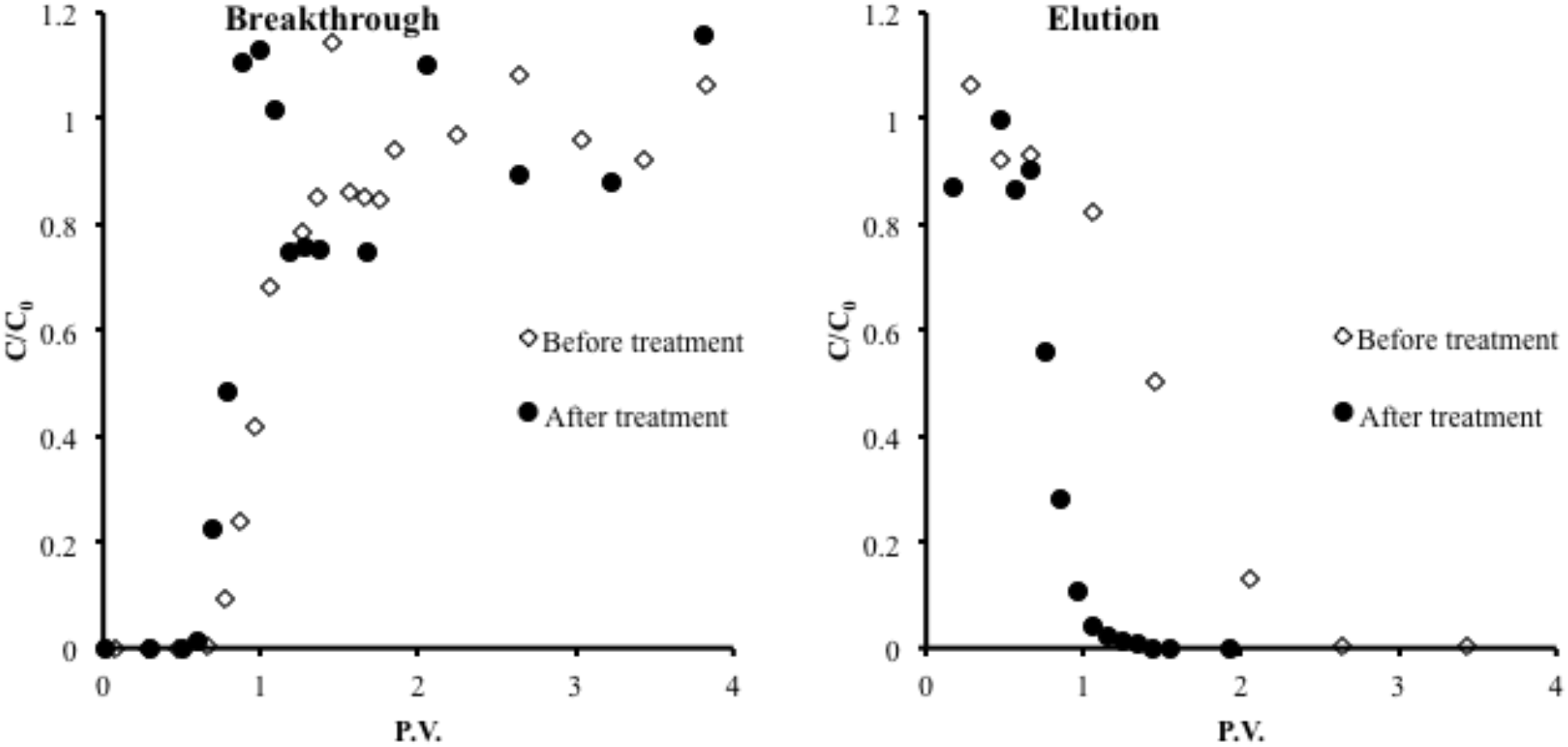
Expanded view of breakthrough and elution profiles of DFBA before and after the 20-day Fe(III)-bioreduction phase.

**Table 2.**
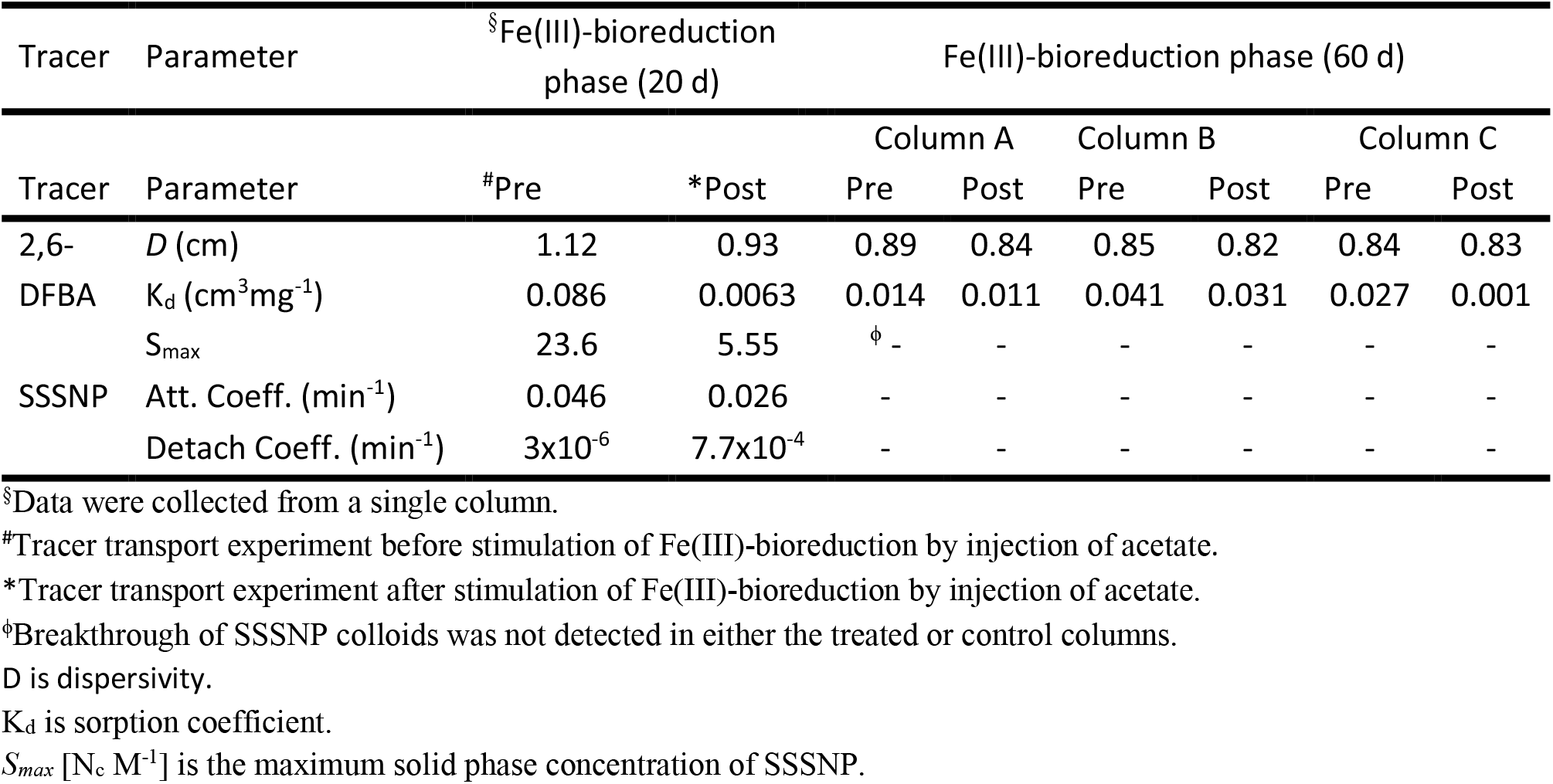
Model fitted parameters for 2,6-DFBA and SSSNP breakthrough from water-stable soil aggregate columns before (Pre-) and after (Post-) Fe(III)-bioreduction.

The detection of soluble Fe(II) in the effluent indicated reduction of soil Fe(III) by *Geobacter* and/or other Fe(III)-reducing bacteria (Fig. 4). The increase in effluent Fe(II) concentrations in the first two weeks corresponded to the increase in microbial activity as acetate was added to the feed solution (Fig. 4). The effluent Fe(II) concentrations plateaued after 20 days of acetate injection, suggesting retardation of Fe(III) bioreduction (Fig. 5). The accumulated amount of Fe(II) in the effluent was approximately 0.125 mmol or about 0.04% of the total iron in the column. No Fe(III) oxides or other colloids were detected in the effluent during the 20-day bioreduction treatment. Although only a very small fraction of total Fe(III) was reduced to soluble Fe(II) (assuming there were no secondary precipitation of Fe(II) in the column), the influences of Fe(III) reduction on the transport of DFBA and SSSNP were significant. This effect is attributable to Fe(II) binding to Fe(III) solids and/or the reduction of the most bioavailable Fe(III) on the external surfaces of the aggregates and/or along the advective pathways that are most accessible by less diffusible microorganisms and particle tracers (e.g., SSSNP). Removal of surface iron oxides during bioreduction could reduce positive electric charges and roughness on mineral surfaces (Joe-Wong et al., 2017), causing a decrease in surface deposition of nanoparticles.

**Fig. 5.**
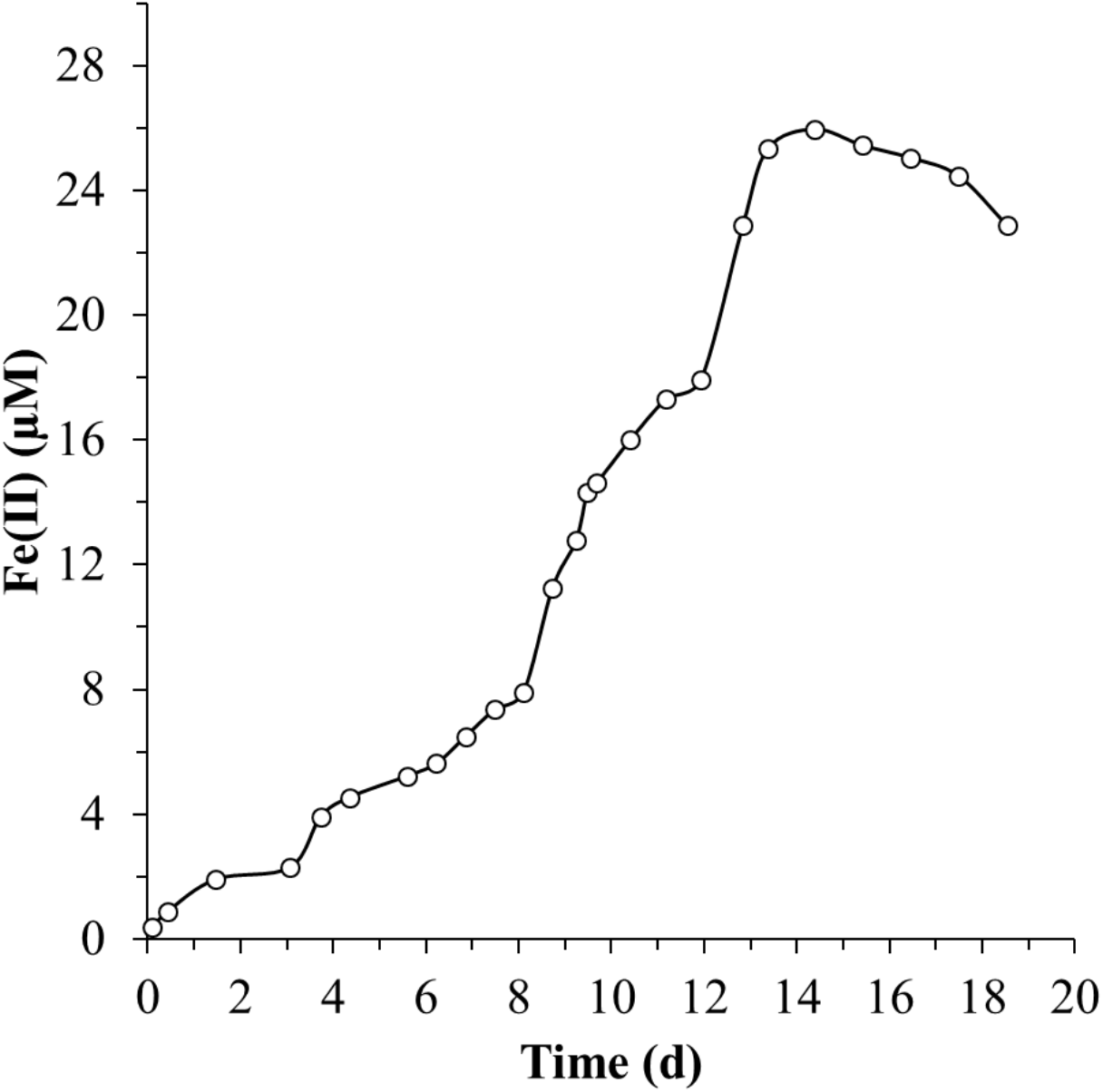
Effluent Fe(II) concentration during the acetate injection phase of the 20-day Fe(III)-bioreduction column experiment.

An in-situ study investigating the mobility of arsenate in natural groundwater showed that arsenic desorption occurred with reductive dissolution of ferric oxides in ferrihydrite, goethite, and hematite (Zhang et al., 2017). Microbial metabolism can affect the affinity of nanoparticles, and therefore cause the simultaneous releases of Fe and Fe(III)-oxide-bound particles. Similarly, Moon et al. (2017) reported that the genus *Geobacter, Anaeromyxobacter*, and *Desulfosporosinus* might play important roles in release of arsenic coupled with iron reduction. In this study, the concentration of Fe(II) in the effluent increased with time during the 20-day treatment demonstrating bioreduction of iron oxides in the acetate-treated soil column. However, without Fe(III) oxides or other colloids detected in the effluent, no evidence of structural breakdown in soil aggregates was observed during the whole bioreduction process. The enhanced transport of SSSNP was most likely attributed to the microbial reduction-induced transformation of iron minerals, resulting in less contact of SSSNP with iron oxides. A very recent study by Xiao et al. (2018) reported that the transformation from less crystalline to more crystalline iron oxides by iron reducing bacterium *Shewanella oneidensis* MR-1 affected the behavior of any species absorbed to the iron oxides, suggesting that the produced Fe(II) can stimulate the reduction and transformation of iron oxide minerals.

### 3.4 Effect of long time bioreduction on transport

Given only 0.04% of the total iron was detected in the effluent in the 20-day bioreduction experiment, additional aggregate-packed column experiments with the duration of acetate-stimulated Fe(III)-bioreduction extended to 60 days were conducted to further examine the effect of bioreduction on Fe(II) release, aggregate breakdown, and tracer transport. The transport behaviors of bromide and DFBA showed no change after the 60-day Fe(III)-bioreduction treatment, whereas SSSNP were completely retained in the soil aggregates both before and after the 60-day acetate injection (Fig. 6 and Table 2). These results are inconsistent with the observations made in the short time experiment with 20-day acetate injection, where Fe(III)-bioreduction resulted in earlier breakthrough of DFBA and SSSNP. The longer duration of Fe(III)-bioreduction (60 d) brought about as yet unknown changes to the properties of soil aggregates and resulted in complete retention of the SSSNP particles, though retardation and slow release of the DFBA were not observed in the short time experiment.

**Fig. 6.**
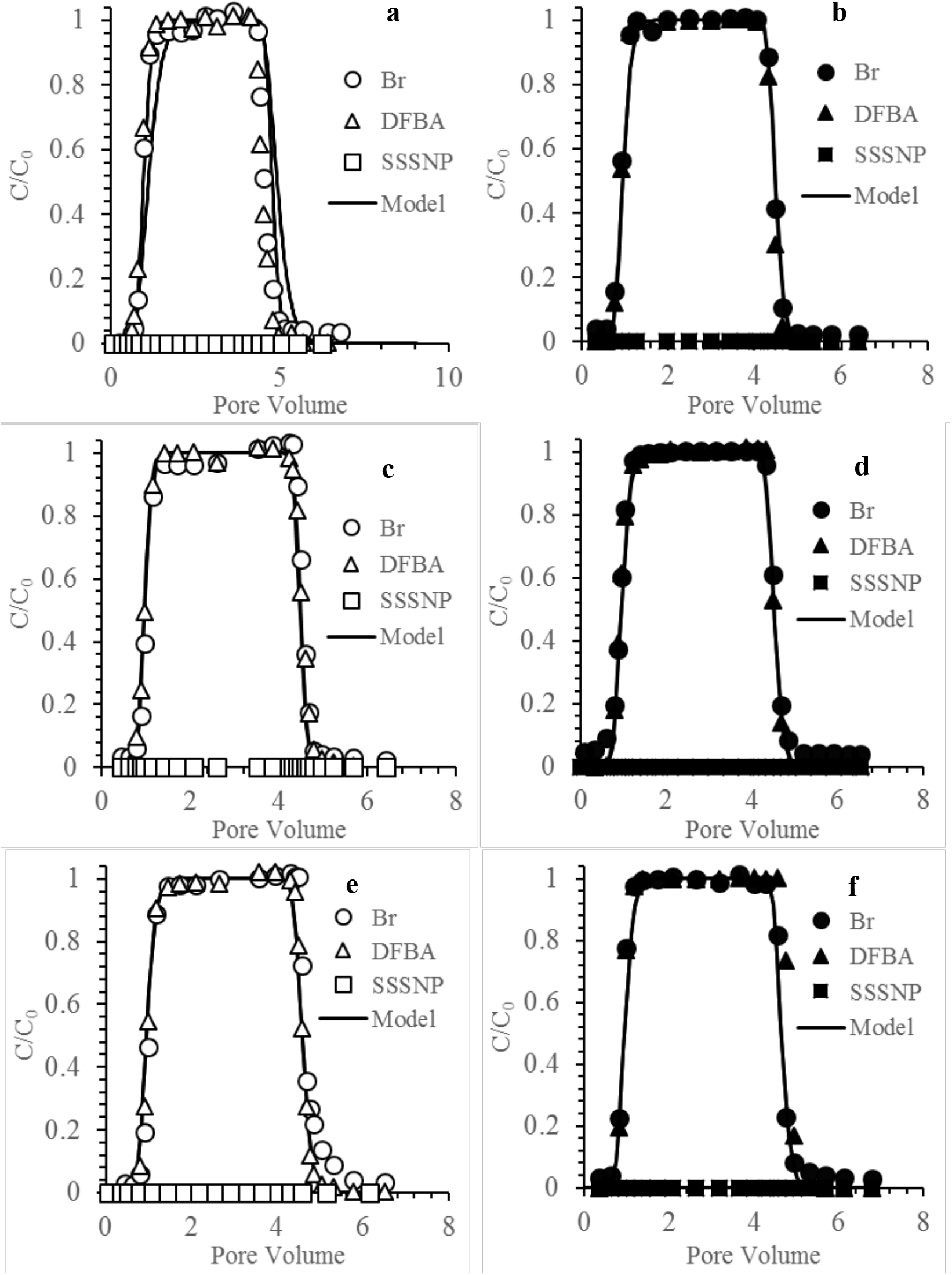
Breakthrough curves of tracers (Br^-^ and DFBA) and silica-shelled silver nanoparticles (SSSNP) in pre-(left, a, c, e) and post-acetate (right, b, d, f) treatment transport experiments for 60-day Fe(III)-bioreduction treatment in the columns packed with water-stable macroaggregates (250-2,000 μm). The plots of a, b, c, and d are breakthrough curves from replicate acetate-treated columns while the plots of e and f are breakthrough curves from the control column that did not receive acetate injection.

The release of soluble Fe(II) in the 60-day experiment was similar to the 20-day experiments during the first 20 days but exhibited a steady increase in the effluent Fe(II) concentration through day 60 (Fig. 7). The concentration of soluble Fe(II) reached 300 μM, or more than 10 times the total amount observed at the end of the 20-day Fe(III) bioreduction experiment. The accumulated amount of soluble Fe(II) collected in the effluent was 1.43 mmol or about 0.48% of the total Fe in the soil aggregates within the column. Colloids were first detected in the effluent at 30 days after the acetate injection and continued to increase in concentration during the 60-day Fe(III)-bioreduction treatment, further suggesting that Fe(III)-bioreduction and aggregate breakdown were active throughout the biostimulation phase of the soil column experiment (Fig. 7). A small amount of Fe(II) was also detected in the effluent of the control column, indicating that *Geobacter* was able to couple oxidation of the native soil organic carbon with Fe(III) reduction. The inconsistent tracer transport results between the 20-day and 60-day bioreduction experiments very likely arose from the greater aggregate breakdown that occurred during the extended period (day 20-60) than the initial period (day 0-20) of the bioreduction treatment. The aggregate breakdown generated soil colloids, which may have increased mechanical straining of SSSNP in soil pores and may have exposed aggregate interior Fe(III)-oxide surfaces promoting the attachment of SSSNP on positively charged surfaces.

**Fig. 7.**
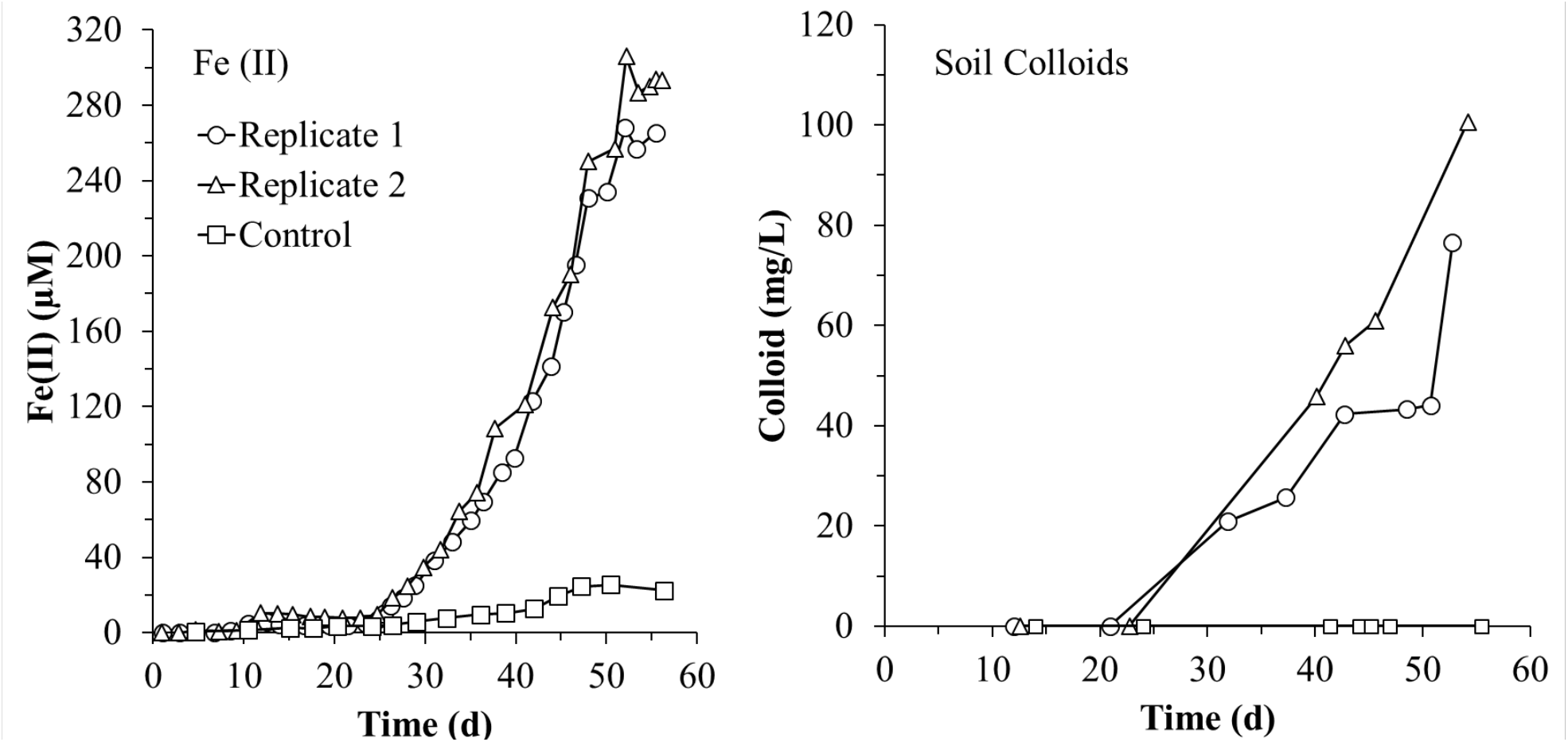
Concentrations of released Fe(II) and soil colloids in the effluent during the acetate injection phase of the 60-day Fe(III)-bioreduction column experiments.

### 3.5 Effect of long time bioreduction on aggregate structure

The above results suggested that the soil aggregates experienced structural breakdown during the 60-day Fe(III)-bioreduction phase with acetate injection. To get direct evidence, we characterized the size distribution of water-stable soil aggregates in each column. The acetate-treated columns contained significantly more soil aggregates with size less than 90 μm than the control column (i.e., no acetate injection) (*P* < 0.05, Fig. 8). The aggregate fractions with sizes of 150-2,000 μm in acetate-treated columns were similar to those in the control column. It is obvious that the soil aggregates enduring 60-day bioreduction generated more microaggregates compared to the soil aggregates with 20-day bioreduction.

**Fig. 8.**
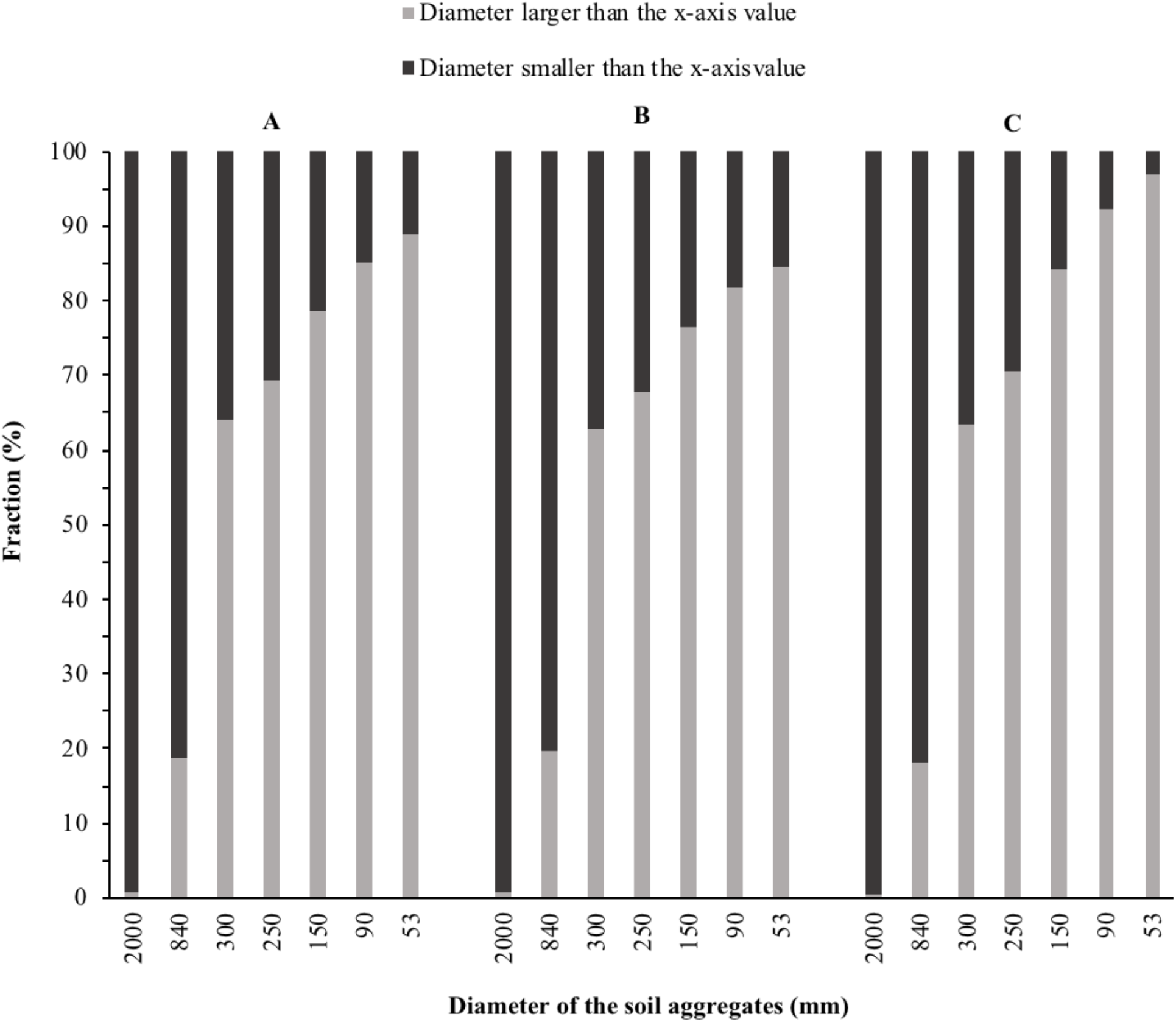
The aggregate size distribution of the soil aggregates from the aggregate-packed columns after the conclusion of the 60-day Fe(III)-bioreduction experiment. Columns A and B received acetate in the feed solution over a 60-day period. Column C was the control column without acetate injection.

Our results also show that the bioreduction increased releases of iron and soil colloids. The divergence of soil aggregates along the length of the columns was also examined by measuring residual water-extractable total Fe. The readily reducible iron contained in the soil aggregates was much higher in the influent sections of the 60-day acetate-fed columns (A and B) with a range of 20-25 mg g^-1^ compared to the control (Fig. 9). The readily reducible iron in the effluent sections of the acetate-fed columns was less than 5 mg g^-1^. In the control column, the water-extractable iron content ranged between 4.5 and 5.5 mg g^-1^ in the influent sections to less than 1 mg g^-1^ in the effluent section. The amounts of readily reducible iron in the acetate-amended columns exceeded that of the control column by approximately five fold. These trends were very consistent with the distribution of the relative abundance of *Geobacter* in the columns (Fig. 9), suggesting that Fe(III)-bioreduction by *Geobacter* contributed to the releases of iron and colloids. These results indicate more significant structural breakdown of the soil aggregates in the soil depths receiving more electron donors.

**Fig. 9.**
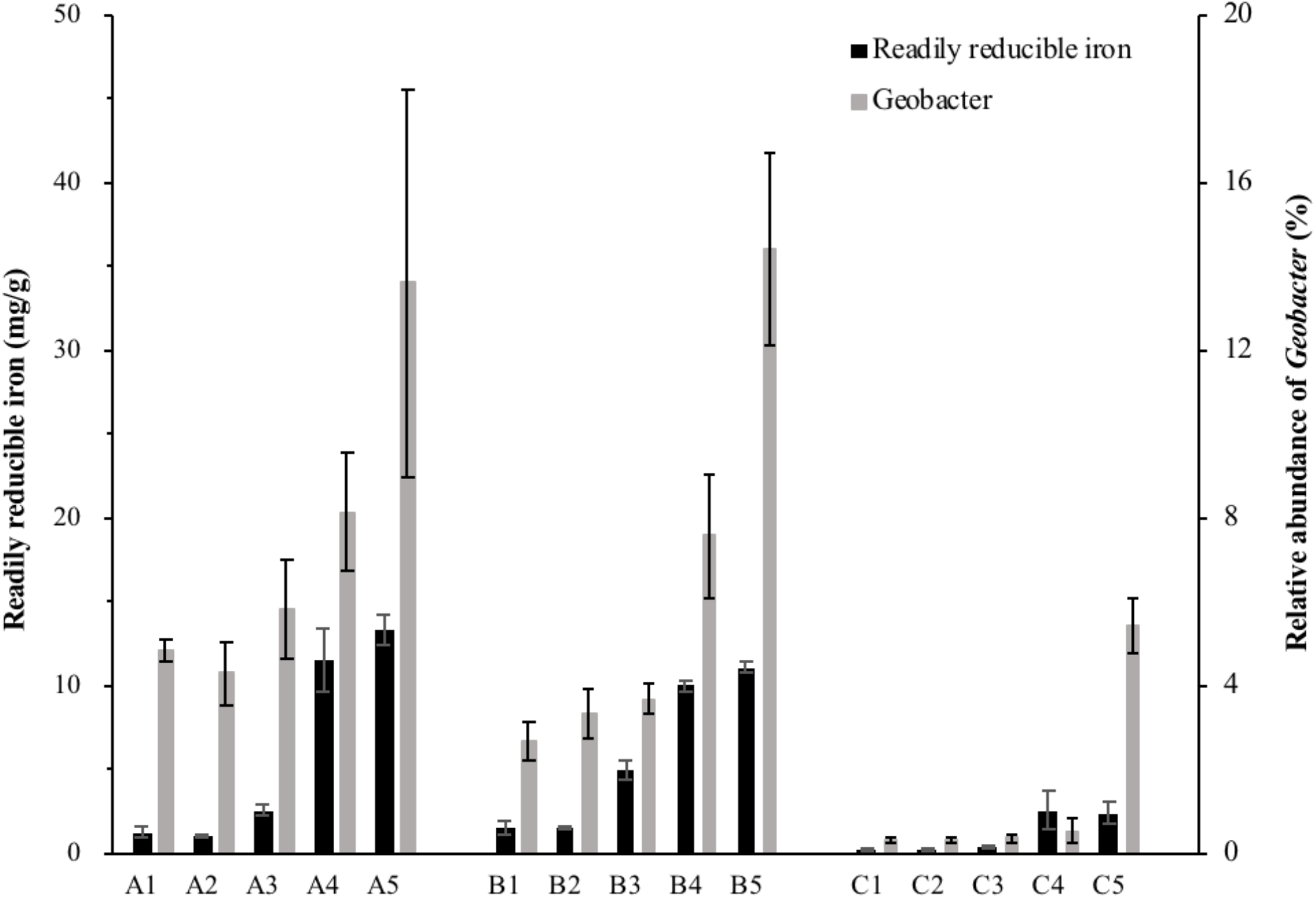
Readily reducible iron content and distribution of iron-reducing bacteria *(Geobacter* determined via sequencing of 16S rRNA gene libraries prepared from soil samples) in five depths of the aggregate-packed columns at the conclusion of the 60-day Fe(III)-bioreduction column experiment. Columns A and B received acetate in the feed solution over a 60-day period. Column C, the control column, did not receive any acetate. Sections 1 and 5 represent samples collected near the effluent and influent ends of the columns, respectively.

## 4. Conclusions and implications

Acetate injection stimulated microbially mediated Fe(III)-oxide reduction, and when delivered for a relatively short duration (i.e. 20 d) it enhanced the transport of an organic molecular tracer (DFBA) and nanoparticle tracers (silica-shelled silver nanoparticles) compared with the transport exhibited prior to acetate injection. However, in the subsequent experiment, when the acetate injection period was extended to 60 d the impact on transport disappeared and the transport of all tracers was identical in acetate treated and control columns despite significant aggregate structural breakdown in the acetate treated columns. The limited data suggest that soil aggregates had minimal structural breakdown during the first 20-day of bioreduction, and as a result, only Fe(III)-oxides coating the exterior surfaces of the aggregates were reduced, yielding advective flow paths with chemically less reactive surfaces that were unfavorable for the attachment of organic and colloidal tracers on the aggregates. In comparison, more aggregates were dispersed during the 60-day bioreduction experiment, causing exposure of interior Fe(III)-oxide surfaces, generating larger reactive surfaces that were favorable for the attachment of organic and colloidal tracers. As a result, the initial 20-day effect was canceled by the subsequent 40-day of extended acetate injection, leading to similar breakthrough behaviors to those observed before the Fe(III)-bioreduction phase. Electron donor addition during biostimulation cannot continue indefinitely. Thus, upon termination of biostimulation, the treated area will ultimately return to its original redox status as oxygenated groundwater passes through the treatment zone. Future studies should address the influence of Fe(II) re-oxidation on tracer transport in relation to changes in soil properties, such as, pore structure and aggregate surface charges along the flow path.

## Acknowledgments

The authors acknowledge with tremendous respect, the contributions of Dr. Phillip Jardine (posthumously) and his collaborators (Drs. Colleen Hansel, Jack Parker, Ungtae Kim, and Kirk Scheckel) for conceiving and executing the research upon which the current work was founded. This research was financially supported by the Strategic Environmental Research and Development Program (SERDP) under project ER-2130. We also acknowledge financial support from the China Scholarship Council to support X.L.

---

**Table S1.**
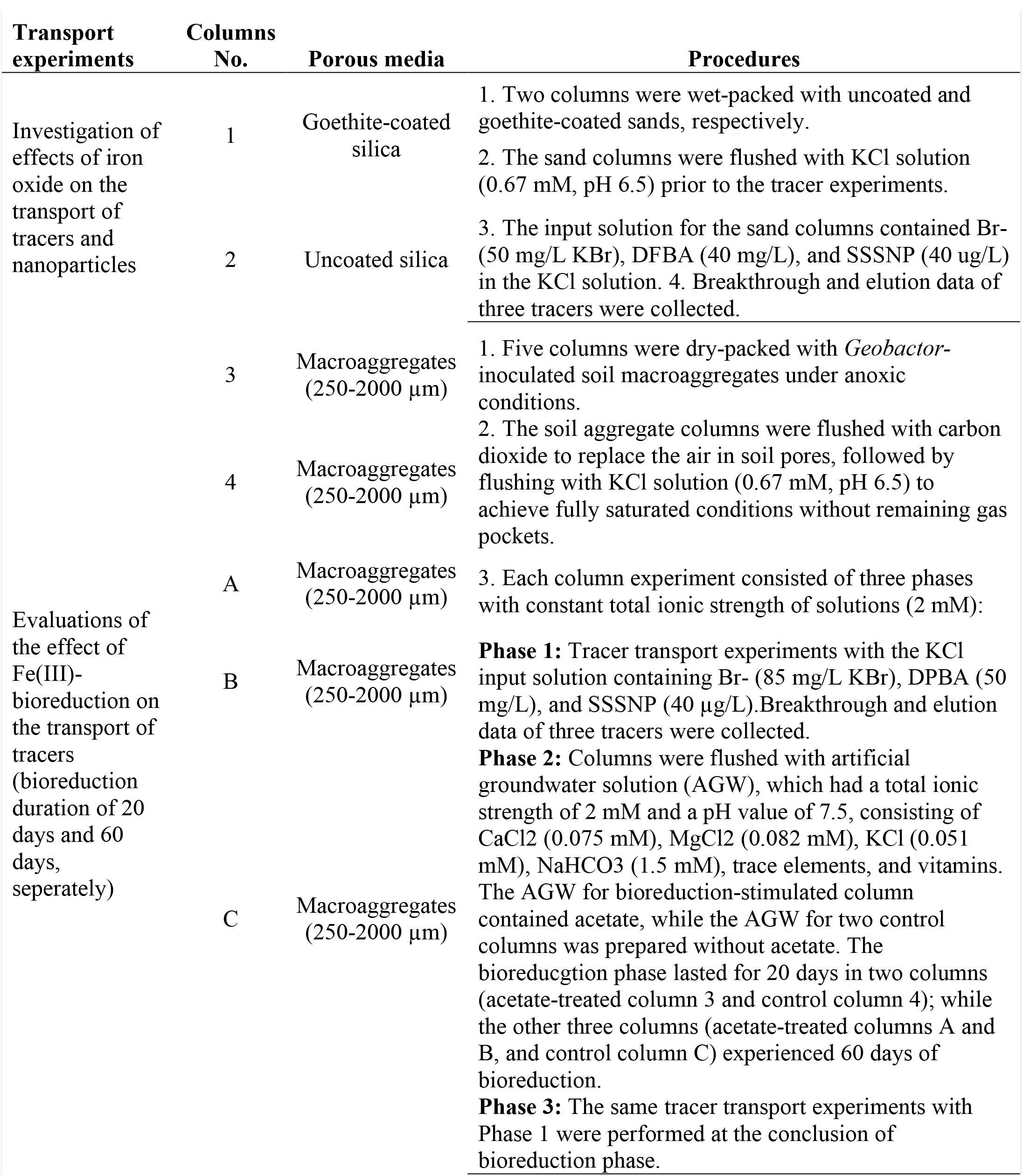
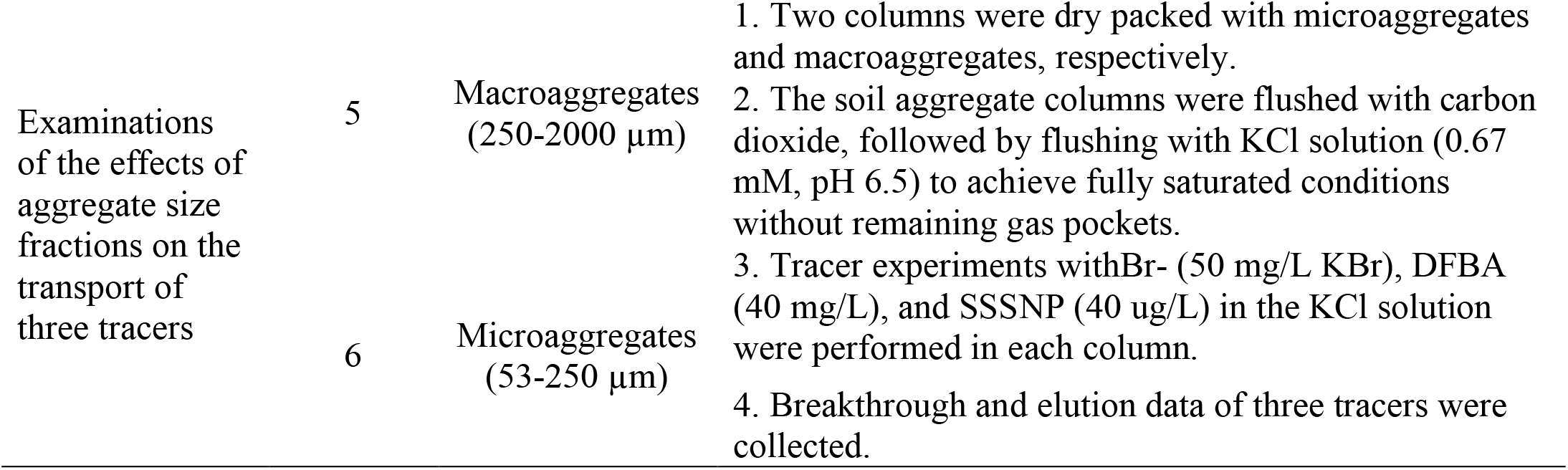
Experimental protocols of transport tests in columns.

**Fig. S1.**
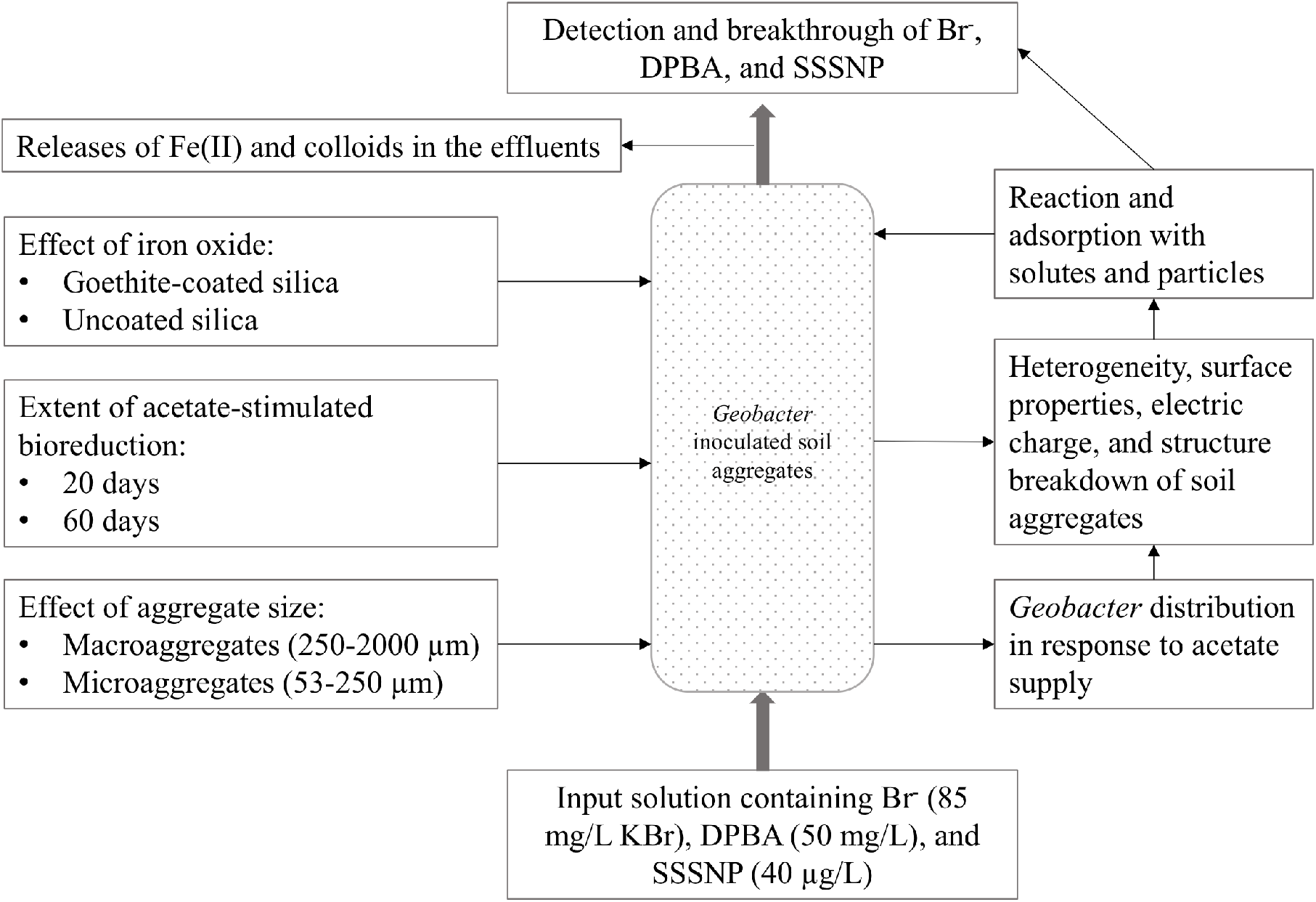
The schematic diagram of column experiments.

